# AVP neurons act as the primary circadian pacesetter cells in vivo

**DOI:** 10.1101/2022.08.04.502742

**Authors:** Yusuke Tsuno, Yubo Peng, Shin-ichi Horike, Kanato Yamagata, Mizuki Sugiyama, Takahiro J. Nakamura, Takiko Daikoku, Takashi Maejima, Michihiro Mieda

## Abstract

The central circadian clock of the suprachiasmatic nucleus (SCN) is a network consisting of various neurons and glia. Individual cells have the autonomous molecular machinery of a cellular clock, but their intrinsic periods are considerably variable. Here, we show that arginine vasopressin (AVP) neurons set the ensemble period of the SCN network to control circadian behavior rhythm. Artificial lengthening of cellular periods by deleting *casein kinase 1 delta* (*CK1δ*) in the whole SCN lengthened the free-running period of behavior rhythm to an extent similar to *CK1δ* deletion specific to AVP neurons. In SCN slices, PER2::LUC reporter rhythms of these mice did not recapitulate the period lengthening. However, in vivo calcium rhythms of both AVP and vasoactive intestinal peptide (VIP) neurons demonstrated lengthened periods similar to the behavioral rhythm upon AVP neuron-specific *CK1δ* deletion. These results indicate that AVP neurons act as the primary determinant of the SCN ensemble period.

## Introduction

In mammals, the suprachiasmatic nucleus (SCN) of the hypothalamus functions as the central clock to control multiple circadian biological rhythms of behaviors and physiological functions, such as sleep-wakefulness, body temperature, and hormone secretion (*1*). Most of approximately 20,000 SCN neurons are GABAergic and include several neuron types characterized by co-expressing peptides. For example, vasoactive intestinal peptide (VIP)-positive neurons in the ventral core region and arginine vasopressin (AVP)-positive neurons in the dorsal shell are two representative neuron types in the SCN (*1*). In individual cells, the molecular machinery of cellular clocks is driven by the autoregulatory transcriptiontranslation feedback loop (TTFL), in which transcriptional activators CLOCK and BMAL1 play a central role (*2*). *Period* (*Per1, 2*, and *3*) and *Cryptochrome* (*Cry1* and *2*) are two of many target genes of CLOCK/BMAL1. PER and CRY proteins then repress CLOCK/BMAL1 activity, completing a negative feedback loop. Casein kinase 1 delta (CK1δ) has been known as a critical regulator of the period length of cellular clocks. By phosphorylating PER2 protein, CK1δ regulates the speed of degradation and nuclear retention of PER2 and, thereby, other clock proteins (*3–6*). Indeed, pharmacological and genetic experiments revealed that eliminating CK1δ activity lengthened periods of circadian behavior rhythm and molecular oscillations in the SCN and peripheral cells (*4, 7, 8*). Intriguingly, these intracellular molecular mechanisms are not unique to SCN neurons but are common to peripheral cells. Instead, intercellular communications among SCN cells are likely essential for the SCN to generate a highly robust, coherent circadian rhythm as the central clock (*1*).

Thus, SCN contains tens of thousands of cellular clocks in various types of neurons and glial cells. However, these cell-autonomous rhythms are sloppy, with a considerable variation in the period length (*1, 9–13*) These facts raise a fundamental question on how the ensemble amplitude, period, and phase of the circadian rhythm at the SCN network level are determined. VIP has been known as the most critical contributor to the synchronization among SCN neurons and is also involved in the photoentrainment to regulate the ensemble phase of the SCN according to the light/dark cycle (*14–20*) On the other hand, recent studies utilizing cell-type-specific genetic manipulations in mice implicated neuromedin S (NMS)-, AVP-, D1a dopamine receptor (DRD1a)-, and VIP receptor VPAC2-expressing neurons and even astrocytes in the pacemaking of the SCN network (*21–27*). NMS neurons were reported to include AVP, VIP, and other types of neurons and act as pacemakers essential for the generation and period-setting of the SCN ensemble rhythm measured by both wheel-running behavior (in vivo) and PER2::LUC reporter expression in slices (ex vivo) (*21*). In contrast, VPAC2 neurons are primarily distributed in the SCN shell and include AVP neurons but not VIP neurons. They regulate the period and coherence of circadian wheel-running rhythm but also require the cooperative action of VIP neurons for pacemaking PER2::LUC rhythms in SCN slices (*25, 28*).

AVP neuron-specific disruption of cellular clocks by deleting *Bmal1* disturbed substantially, but not completely abolished, the circadian rhythm of locomotor activity (home-cage activity) (*22*). Also, manipulating the cellular circadian period of AVP neurons by deleting (lengthening) or overexpressing (shortening) *CK1δ* altered the free-running period of locomotor activity rhythm accordingly (*23*). These observations support a significant role for AVP neurons in the period-setting and coherence of the SCN ensemble rhythm. However, the behavioral period lengthening by the specific deletion of *CK1δ* was not fully recapitulated in cellular PER2::LUC rhythms in SCN slices (*23*), implicating different states of the SCN network between in vivo and ex vivo. In addition, the extent to which AVP neurons contribute to the SCN pacemaking relative to other types of SCN neurons remains unclear. To address these questions, we compared the period-lengthening effects caused by *CK1δ* deletion in the entire SCN to AVP neuron-specific deletion in both behavior and slice rhythms. In addition, we examined whether the behavioral period lengthening due to AVP neuron-specific *CK1δ* deletion is recapitulated in the SCN cellular clock oscillations in vivo by monitoring the intracellular Ca^2+^ ([Ca^2+^]_i_) rhythms of AVP and VIP neurons using fiber photometry in freely behaving animals. Our results suggested that AVP neurons act as the principal pacesetter cells in the SCN network in vivo.

## Results

### Deletion of *CK1δ* in the entire SCN lengthens the circadian period of behavior comparably to AVP neuron-specific *CK1δ* deletion

We previously showed that lengthening the cellular circadian period of AVP neurons by the specific deletion of *CK1δ* lengthened the free-running period of circadian behavior, indicating that AVP neurons are involved in setting the ensemble period of the SCN network (*23*). The remaining question was how much AVP neurons contribute to the period setting. To elucidate this, we aimed to compare the period lengthening caused by AVP neuron-specific *CK1δ* deletion to that by pan-SCN *CK1δ* deletion. Although CaMKIIα is not abundant in the SCN (*29*), a particular *CaMKIIα-Cre* line (*30*) shows clear expression not only in the forebrain but also in the SCN (*31*). Indeed, crossing this line to floxed *Bmal1* mice resulted in >90% deletion of BMAL1 in the SCN and a complete loss of circadian behavior rhythm (*31*). We also reproduced the arrhythmicity of *CaMKIIα-Cre; BmalKl1^flox/flox^* mice, confirming *CaMKIIα-Cre* mice as a pan-SCN Cre driver (Sup. Fig. S1A-B).

By crossing this *CaMKIIα-Cre* line with *CK1δ^flox/flox^* (*8*), we deleted *CK1δ* in the forebrain and the entire SCN neurons (*CaMKIIα-CK1δ^-/-^*). *CaMKIIα-CK1δ^-/-^* mice showed normal diurnal locomotor activity (home-cage activity) rhythm in the 12 h of light and 12 h of darkness (LD) condition (Fig. 1A, 1B). In constant darkness (DD), *CaMKIIα-CK1δ^-/-^* mice showed a longer free-running period (24.83 ± 0.08 h) than that of control mice (23.91 ± 0.03 h, *P* < 0.001, Fig. 1C). By contrast, their amplitude of locomotor activity rhythm (Qp) was normal (*P* = 0.53, Fig. 1D). The difference in the free-running period between control and *CaMKIIα-CK1δ^-/-^* mice was almost similar to the difference between control and *Avp-CK1δ^-/-^* mice (23.94 ± 0.03 h vs. 24.72 ± 0.03 h) (*23*), as well as with that between control and *Vgat-Cre; CK1δ^flox/flox^* mice (approximately 40 min elongation), another line with pan-SCN neuronal *CK1δ* deficiency (*32*). These data suggest that AVP neurons are the principal determinant of the free-running period of the circadian behavior rhythm.

**Figure 1.**
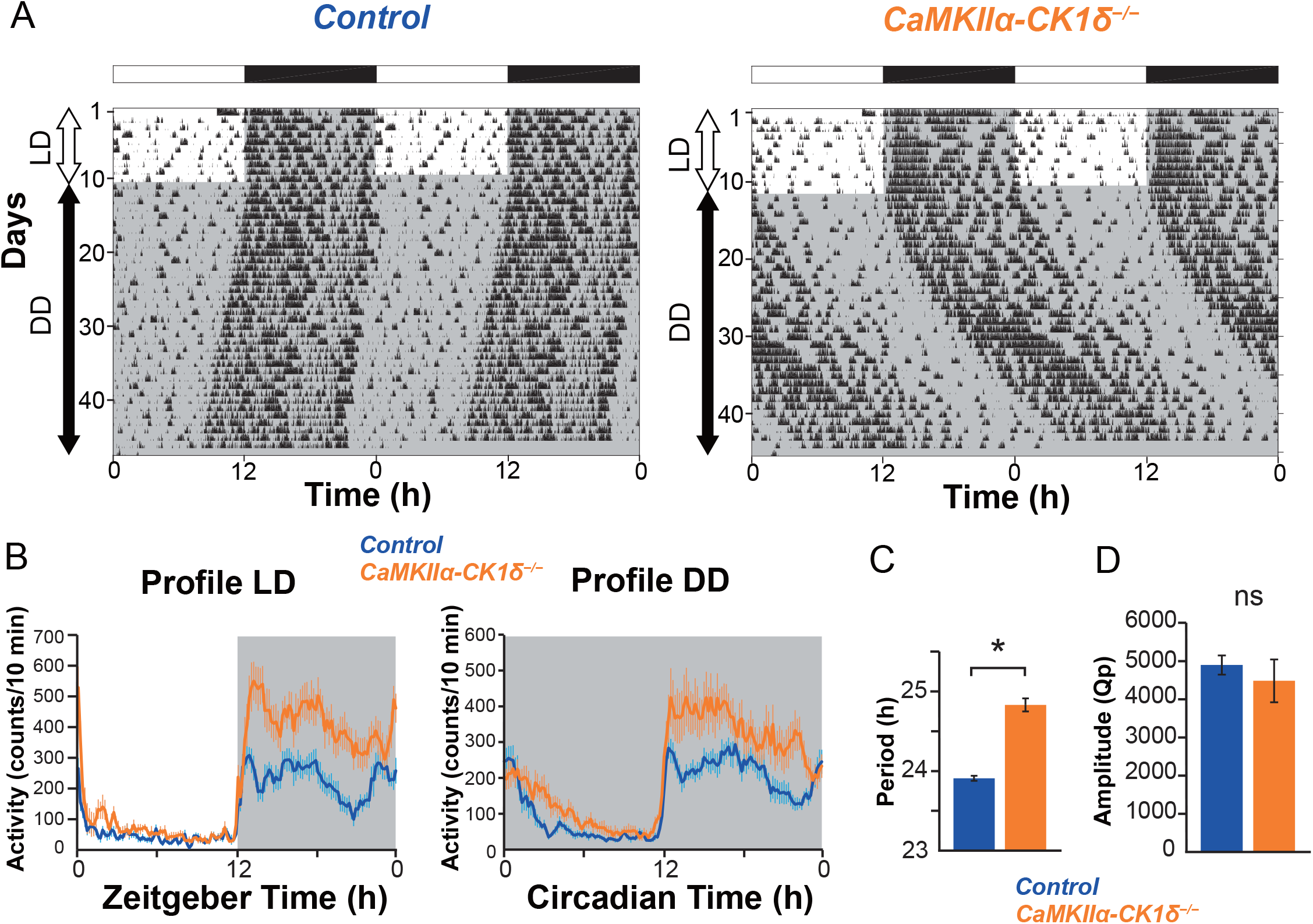
*CaMKIIa-CK1δ^-/-^* mice show lengthening of the free-running period in DD. (A) Representative locomotor activity of control and *CaMKIIa-CK1δ^-/-^* mice. Animals were initially housed in 12:12-h LD conditions and then transferred to DD. Gray shading indicates the time when lights were off. (B) Averaged daily profile of locomotor activity in LD (left) or DD (right). (C) The free-running period in DD. (D) The circadian amplitude of locomotor activity rhythms (Qp values obtained from periodogram analyses).Values are mean ± SEM; n = 11 for control, n = 13 for *CaMKIIa-CK1δ^-/-^* mice. *\*P* < 0.05 by two-tailed Student’ s t tests; ns, not significant.

To examine the contribution of VIP neurons in the setting of the SCN ensemble period, we deleted *CK1δ* specifically in VIP neurons by crossing *Vip-ires-Cre* mice (*33*) with *CK1δ^flox/flox^* (*Vip-CK1δ^-/-^*). However, we did not observe any change in locomotor activity rhythm of *Vip-CK1δ^-/-^* mice (Fig. 2A-D, period, control 23.93 ± 0.04 vs. *Vip-CK1δ^-/-^* 23.91 ± 0.03). This result was consistent with a previous report that the lengthening of the cellular period by the overexpression of *Clock^Δ19^* did not alter the behavioral free-running period (*34*). These observations suggest that VIP neurons have little contribution to the period setting of the SCN network, although VIP peptide has been established as an essential synchronizer of SCN neurons.

**Figure 2.**
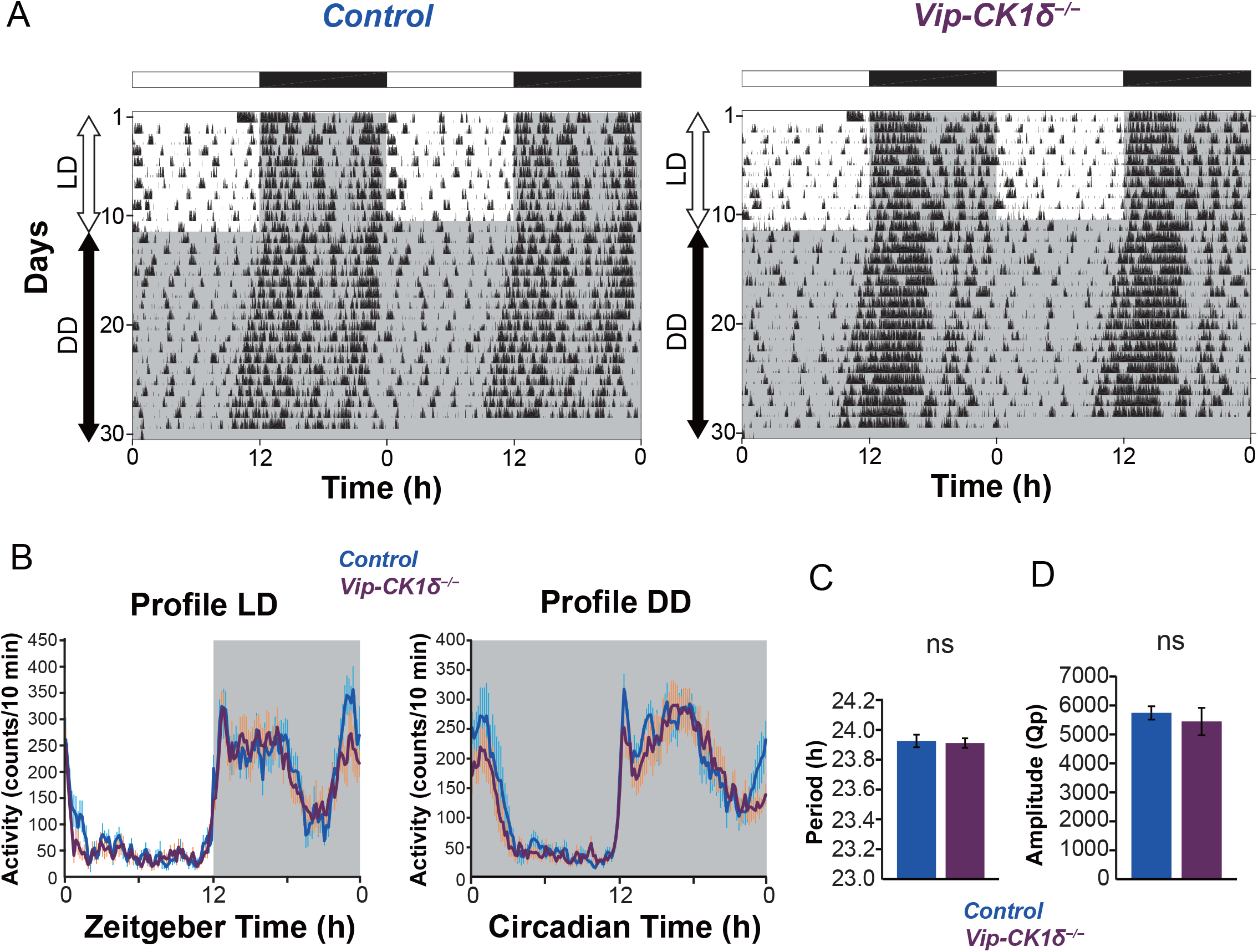
*Vip-CK1δ^-/-^* mice show no change in the free-running period in DD. (A) Representative locomotor activity of control and *Vip-CK1δ^-/-^* mice. Gray shading indicates the time when lights were off. (B) Averaged daily profile of locomotor activity in LD (left) or DD (right). (C) The free-running period in DD. (D) The circadian amplitude of locomotor activity rhythms (Qp values). Values are mean ± SEM; n = 9 for control, n = 9 for *Vip-CK1δ^-/-^* mice. ns, not significant.

### Disrupting cellular clocks in AVP neurons causes arrhythmic wheel-running

We previously reported that disruption of cellular clocks in AVP neurons by deleting *Bmal1 (Avp-Bmal1^-/-^* mice) causes severe attenuation of circadian behavior rhythm measured by home-cage activity (*22*). However, most *Avp-Bmal1^-/-^* mice still retained some circadian rhythmicity with unstable, lengthened free-running period and activity time. In contrast, Lee et al. reported that mice with *Bmal1* deletion in NMS neurons (*Nms-Bmal1^-/-^* mice) were arrhythmic in wheel-running behavior (*34*). Notably, a recent study showed that rhythms of home-cage activity and body temperature in *Avp-Bmal1^-/-^* mice were very similar to those in *Nms-Bmal1^-/-^* mice (*35*). Given the critical role of SCN AVP neurons in setting the free-running period, we re-evaluated the circadian behavior rhythm of *Avp-Bmal1^-/-^* and *Avp-CK1δ^-/-^* mice by measuring wheel-running activity. The difference in wheel-running rhythm between *Avp-CK1δ^-/-^* and control mice was similar to that of home-cage activity rhythm, namely, ~50 min lengthening of the free-running period (control 23.73 ± 0.05 vs. *Avp-CKlδ^-/-^* 24.47 ± 0.07, *P* < 0.001, Sup. Fig. S2, A and B). Strikingly, 7 out of 8 *Avp-Bmal1^-/-^*mice exhibited almost arrhythmic wheel-running (Sup. Fig. S2C). These data suggested that the impairment of circadian behavior rhythm is comparable between *Avp-Bmal1^-/-^* and *Nms-Bmal1^-/-^* mice.

### PER2::LUC oscillations in SCN slices fail to recapitulate lengthened behavioral periods of *CaMKIIα-CK1δ^-/-^* and *Avp-CKlδ^-/-^* mice

To evaluate the status of cellular circadian clocks in the SCN, we next performed real-time bioluminescent cell imaging of coronal SCN slices prepared from control, *Avp-CKlδ^-/-^*, *CaMKIIα-CK1δ^-/-^*, and *Vip-CKlδ^-/-^* adult mice crossed with a luciferase reporter (*Per2::Luc*) housed in LD. In the previous study, we reported that PER2::LUC oscillations in the SCN slices of *Avp-CK1δ^-/-^* mice did not demonstrate coherent circadian rhythm with a lengthened period, contrary to their locomotor activity rhythm (*23*). The SCN shell and core of *Avp-CK1δ^-/-^; Per2::Luc* mice transiently showed different cellular periods ex vivo. The period of the shell was longer by ~2 h at the beginning. However, this lengthening did not continue into the next cycle. In contrast, PER2::LUC expression in the core oscillated stably with a period of ~24 h.

Therefore, we next monitored and compared the day-by-day change of individual pixels’ periods of PER2::LUC oscillations between shell and core, as well as among genotypes. To do so, we calculated the daily peak phase and period (the interval of two adjacent peaks) for PER2::LUC oscillations in individual pixels that covered the SCN (Fig. 3, 4 and Sup. Fig. S3). The results of *Avp-CK1δ^-/-^; Per2::Luc* mice were similar to our previous report. The peak phases in the shell pixels were later than those in core pixels by ~5 h at the first peak (Fig. 3, 4D-F). Also, the periods were longer in the shell (25.77 ± 0.21 h) than in the core (23.68 ± 0.12 h) by ~2 h for the first cycle (Fig. 3 and 4A). While core pixels oscillated with relatively stable periods, those of shell pixels shortened rapidly in the subsequent cycles, as if core cells influenced the periods of shell cells (Fig. 4A and Sup. Fig. S3). Accordingly, the peak phase difference between shell and core did not further increase at the third and fourth peaks (Fig. 4F).

**Figure 3.**
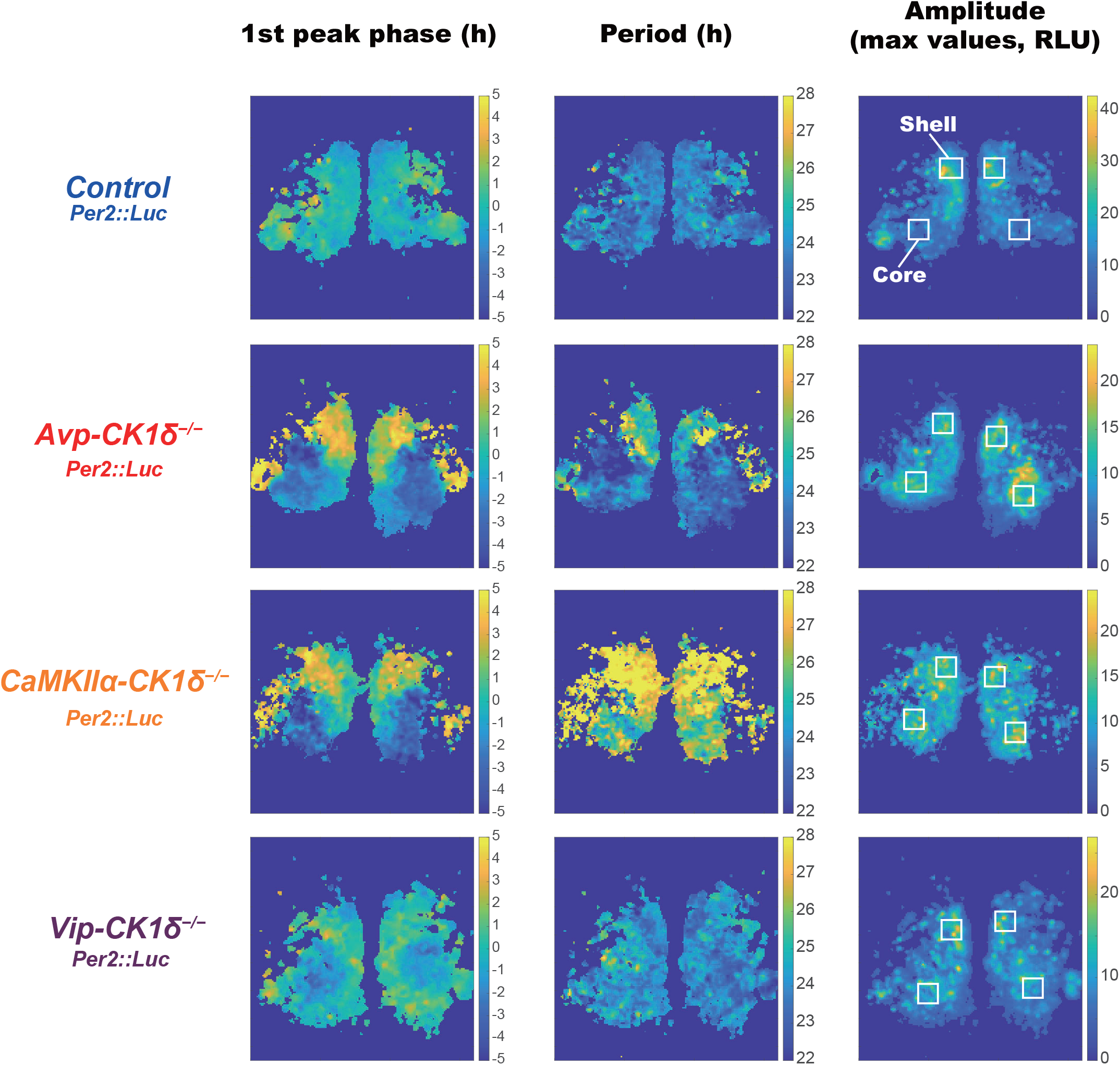
The shell and core transiently show different periods of PER2::LUC oscillation in the SCN slices of *Avp-CK1δ^-/-^* and *CaMKIIa-CK1δ^-/-^* mice. Representative first peak phase (relative phase to the slice mean), period (the interval between the first and second peaks), and amplitude maps of PER2::LUC oscillation at the pixel level in coronal SCN slices prepared from control, *Avp-CK1δ^-/-^, CaMKIIa-CK1δ^-/-^* and *Vip-CK1δ^-/-^* mice. Peak phases, periods, and amplitudes of PER2::LUC oscillations in the individual pixels covering the SCN were calculated for every cycle. White squares on the amplitude maps indicate regions (15 x 15 pixels) considered as the shell or core for further analyses in Fig. 4.

**Figure 4.**
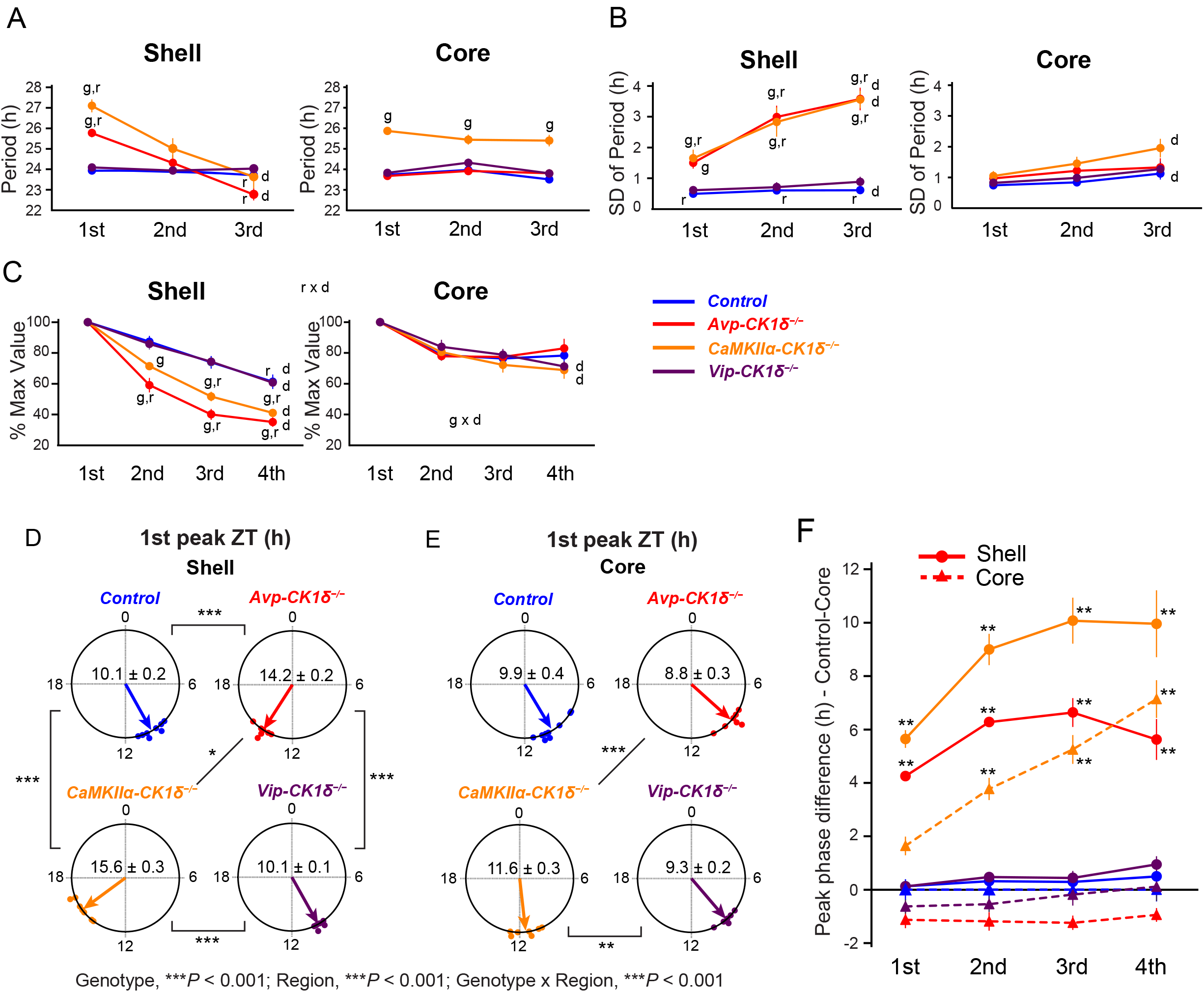
Periods of PER2::LUC oscillations in the SCN shell rapidly changes in the SCN slices of *Avp-CK1δ^-/-^* and *CaMKIIα-CK1δ^-/-^*mice. (A) The day-by-day change in the mean period of individual pixels’ PER2::LUC oscillations in the shell (left) and core (right) regions. The analyzed regions’ examples are shown in Fig. 3. Definitions of peaks, periods, and cycles are indicated in Sup. Fig. S3. The averaged values of two regions in the left and right SCN of individual slices were considered the representatives of individual mice. Blue, Control; Red, *Avp-CKIδ^-/-^* Orange, *CaMKIIα-CKIδ^-/-^* Purple, *Vip-CK1δ^-/-^*. (B) The day-by-day change in the standard deviation (SD) of individual pixels’ periods in the shell (left) and core (right) regions. (C) The day-by-day change in the mean peak amplitude (relative to the 1st peak amplitude) of individual pixels’ PER2::LUC oscillations in the shell (left) and core (right) regions. (D, E) The mean first peak phase (ZT) of the shell (D) and core (E) regions in different mouse lines were shown as Rayleigh plots. Individual dots indicate the mean peak phases of each mouse. (F) Peak phase differences between the SCN core of control mice and the shell or core of other mouse lines. Circle, shell; triangles, core. Values are mean ± SEM; n = 9 for Control, n = 7 for *Avp-CK1δ^-/-^*, n = 7 for *CaMKIIα-CK1δ^-/-^*, n = 6 for *Vip-CK1δ^-/-^*. Letters indicate significant differences in the factor of genotype (g: compared with Control), day (d), or region (r). g: effect of genotype, ***P*** < 0.05 by Kruskal-Wallis rank sum test followed by Man-Whitney U test with Bonferroni correction (A, B) or three-way repeated measures ANOVA with post-hoc Ryan’ s test (C); d: effect of day, ***P*** < 0.05 by Friedman rank sum test followed by Wilcoxon signed rank test with Bonferroni correction (A, B) or three-way repeated measures ANOVA with post-hoc Ryan’ s test (C); r: effect of region, ***P*** < 0.05 by Wilcoxon signed rank test (A, B) or three-way repeated measures ANOVA with post-hoc Ryan’ s test (C); g x d, interaction between genotype and day (C, Core); r x d, interaction between region and day (C, in all genotypes). \****P*** < 0.05; \*\****P*** < 0.01; \*\*\****P*** < 0.001 by Harrison-Kanji test followed by Watson-Williams test with Bonferroni correction (D) or Kruskal-Wallis rank sum test followed by Man-Whitney U test with Bonferroni correction (F, effect of Genotype-Region compared with Control-Core). ***P*** values of Rayleigh test were < 0.01 for all circular data.

In *CaMKIIa-CK1δ^-/-^; Per2::Luc* mice, all SCN neurons appear to have similarly lengthened periods of PER2::LUC rhythms. Therefore, we expected that SCN explants of those mice would show coherent PER2::LUC oscillations similar to those of control mice, except for the lengthening of periods. Indeed, core pixels oscillated stably with lengthened periods (~25 h) for three cycles (Fig. 4A right). However, we observed a rapid temporal change of periods in the shell pixels similar to that in *Avp-CKlδ^-/-^; Per2::Luc* mice, except for ~1 h lengthening (Fig. 4A left). Namely, the peak phases in the shell pixels were later than those in core pixels by ~4 h at the first peak (Fig. 3, 4D–F). In addition, the periods were longer in the shell (27.09 ± 0.32 h) than in the core (25.87 ± 0.11 h) by ~2 h for the first cycle. However, periods of shell pixels shortened rapidly in the subsequent cycles (Fig. 4A). Thus, the coherent circadian rhythm was not maintained in PER2::LUC expression of the SCN slice cultures in *CaMKIIα-CK1δ^-/-^* mice, a common phenomenon with *Avp-CK1δ^-/-^* mice. Concordantly, the variability of the individual pixels’ periods increased in the SCN shell of *Avp-CK1δ^-/-^* and *CaMKIIα-CK1δ^-/-^* mice (Fig. 4B). Furthermore, the peak amplitude of individual pixels’ PER2::LUC oscillations decayed more rapidly in the shell of these two strains of mice (Fig. 4C). These observations suggested the attenuated synchronization of cellular PER2::LUC oscillations in the SCN shell of *Avp-CK1δ^-/-^* and *CaMKIIα-CK1δ^-/-^* mice ex vivo. In contrast, PER2::LUC oscillations in the SCN slices of *Vip-CK1δ^-/-^; Per2::Luc* mice demonstrated no significant difference compared to control mice. This observation was consistent with their normal free-running period of circadian locomotor activity rhythm.

### The circadian period of calcium rhythm in SCN AVP neurons of *Avp-CK1δ^-/-^* mice is stably lengthened in vivo

Lengthening the cellular circadian periods of AVP neurons could stably lengthen the free-running period of the behavioral rhythm. However, it could not lengthen the cellular PER2::LUC rhythms in the prolonged SCN cultures. Therefore, we postulated that the action of AVP neurons might be susceptible to slicing. To test this possibility, we next measured the cellular period of SCN AVP neurons in vivo by recording the intracellular Ca^2+^ ([Ca^2+^]_i_) rhythm using fiber photometry (*36*) in *Avp-CK1δ^-/-^* and control (*Avp-Cre; CK1δ^wt/flox^*) mice (Fig. 5). To do so, we expressed a fluorescent Ca^2+^ indicator jGCaMP7s (*37*) specifically in SCN AVP neurons by focally injecting a Cre-dependent AAV vector (*AAV-CAG-DIO-jGCaMP7s*), then implanted an optical fiber just above the SCN (Fig. 5, A and B). The detected jGCaMP7s signal was averaged, detrended, and smoothened to extract daily [Ca^2+^]_i_ rhythms (see Materials and Methods). We defined the time points crossing zero values as the onset and offset of the GCaMP signal and their midpoint as the peak phase. In both control and *Avp-CK1δ^-/-^* mice, daily [Ca^2+^]_i_ rhythms were observed in SCN AVP neurons in both LD and DD. Their peaks were around the offset of locomotor activity (Fig. 5, C and D, Fig. 7A, and Sup. Fig. S4A), as described previously for control mice (*36*). The relationship between the GCaMP peak and locomotor offset remained unchanged in *Avp-CK1δ^-/-^* mice. Thus, not only the period of locomotor activity rhythm but also the period of [Ca^2+^]_i_ rhythm in AVP neurons was lengthened similarly in DD (GCaMP, control 23.77 ± 0.18 vs. *Avp-CK1δ^-/-^* 24.42 ± 0.13, *P* < 0.05, Fig 7D; behavior, control 23.77 ± 0.10 vs. *Avp-CK1δ^-/-^* 24.43 ± 0.09, *P* < 0.001, Fig. 7F black). This result indicated that the *CK1δ* deletion in AVP neurons caused a stable lengthening of their cellular circadian period in vivo.

**Figure 5.**
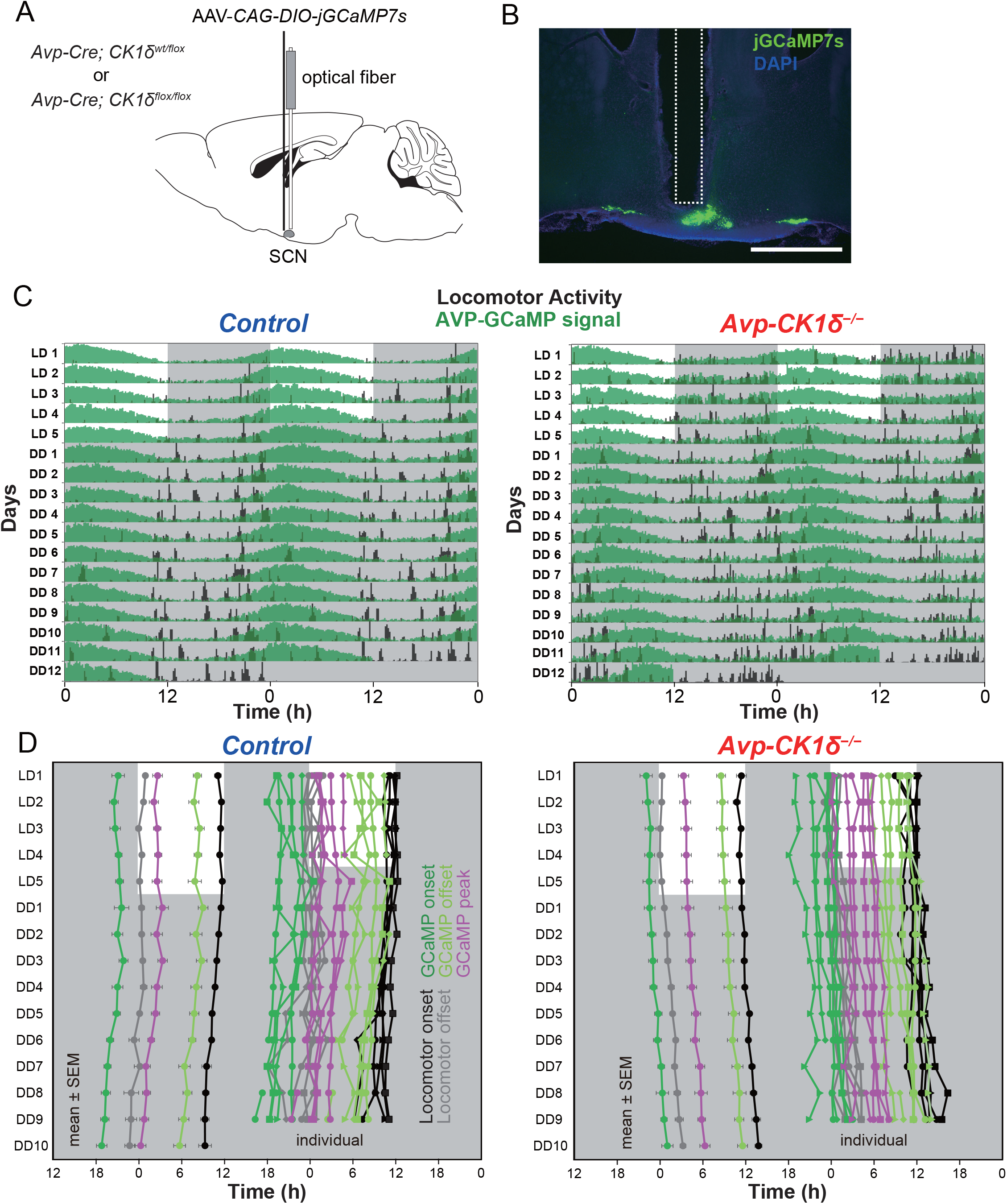
The in vivo circadian period of AVP-neuronal [Ca^2+^] rhythm is lengthened in the SCN of *Avp-CK1δ^-/-^*mice. (A) Schematic diagram of virus vector (*AAV-CAG-DIO-jGCaMP7s*) injection and optical fiber implantation at SCN in control (*Avp-Cre; CK1δ^wt/flox^*) or *Avp-CK1δ^-/-^ (Avp-Cre; CK1δ^flox/flox^*) mice for fiber photometry recording. (B) A representative coronal section of mice with jGCaMP7s expression in SCN AVP neurons. A white dotted square shows the estimated position of implanted optical fiber. Green, jGCaMP7s; blue, DAPI. Scale bar, 1 mm. (C) Representative plots of the in vivo jGCaMP7s signal of SCN AVP neurons (green) overlaid with locomotor activity (black) in actograms. Control (Left) and *Avp-CK1δ^-/-^* (Right) mice were initially housed in LD (LD1 to LD5), then in DD (DD1 to DD12). The dark periods are represented as gray shaded areas. (D) Plots of locomotor activity onset (black), activity offset (gray), GCaMP onset (green), GCaMP offset (light green), and GCaMP peak (magenta) of mean ± SEM (left column) and individual mice data (right column) in control and *Avp-CK1δ^-/-^* mice. Identical marker shapes indicate data from the same animal. n = 5 for control (n = 3 without *Vip-tTA;* n = 2 with *Vip-tTA*), n = 6 for *Avp-CK1δ^-/-^* (n = 4 without *Vip-tTA;* n = 2 with *Vip-tTA*).

### AVP neurons entrain VIP neurons in vivo

For setting the SCN network ensemble period, AVP neurons need to regulate the circadian oscillations of other SCN neurons. Because of the importance of VIP neurons in the SCN core for maintaining the coherence of circadian oscillators in the SCN (*14, 15, 18, 38, 39*), we next examined whether the [Ca^2+^]_i_ rhythm of SCN VIP neurons was altered in *Avp-CK1δ^-/-^* mice in vivo. To specifically target the jGCaMP7s expression in VIP neurons of *Avp-CK1δ^-/-^* mice, we first crossed *Avp-CK1δ^-/-^* (or control) with *Vip-tTA* knock-in mice (*40*). In *Vip-tTA* mice, VIP neurons specifically express tetracycline transactivator (tTA). We then injected a tTA-dependent AAV vector (AAV*-TRE-jGCaMP7s*) into the SCN (Fig. 6, A and B). The [Ca^2+^]_i_ rhythms of VIP neurons were synchronized antiphasically with locomotor activity rhythm in both control (*Avp-Cre;CK1δ^wt/flox^; Vip-tTA*) and *Avp-CK1δ^-/-^; Vip-tTA* mice (Fig. 6, C and D, Sup. Fig. S4B), as reported previously for control mice (*18, 36, 41*) Furthermore, the timing of locomotor onset and GCaMP offset were almost identical, while the timing of locomotor offset and VIP GCaMP onset correlated in both LD and DD (Fig. 6, C and D). In *Avp-CK1δ^-/-^* mice, the period of VIP-neuronal [Ca^2+^]_i_ rhythm was lengthened in DD to the same extent as AVP-neuronal [Ca^2+^]_i_ and behavior rhythms (GCaMP, control 23.87 ± 0.03 vs. *Avp-CK1δ^-/-^* 24.47 ± 0.05, *P* < 0.001, Fig. 7E). Importantly, as a control, we verified that EGFP-expressing VIP neurons via injecting AAV*-TRE-EGFP* showed very little circadian oscillation of fluorescence (Sup. Fig. S4C). These results suggested that the cellular clocks of AVP neurons could regulate those of VIP neurons.

**Figure 6.**
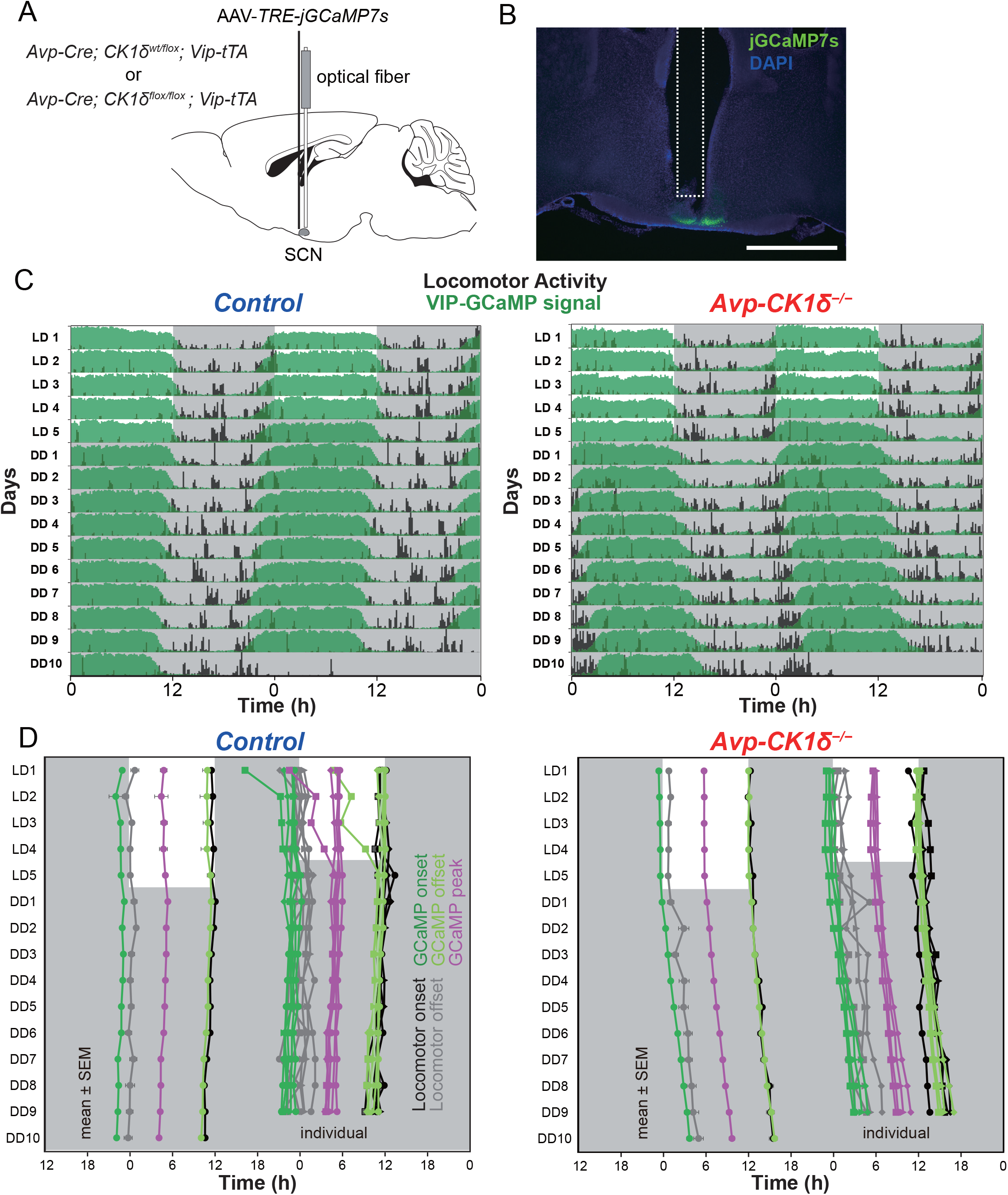
The in vivo cirnadian period of VIP-neuronal [Ca^2+^]_i_ rhythm is lengthened in the SCN of *Avp-CK1δ^-/-^*mice. (A) Schematic diagram of virus vector (*AAV-TRE-jGCaMP7s*) injection and optical fiber implantation at SCN in control (*Avp-Cre; CK1δ^wt/flox^; Vip-tTA*) or *Avp-CK1δ^-/-^ (Avp-Cre; CK1δ^wt/flox^; Vip-tTA*) mice for fiber photometry recording. (B) A representative coronal section of mice with jGCaMP7s expression in SCN VIP neurons. A white dotted square shows the estimated position of implanted optical fiber. Green, jGCaMP7s; blue, DAPI. Scale bar, 1 mm. (C) Representative plots of the in vivo jGCaMP7s signal of SCN VIP neurons (green) overlaid with locomotor activity (black) in actograms. Control (Left) and *Avp-CK1δ^-/-^* (Right) mice were initially housed in LD (LD1 to LD5), then in DD (DD1 to DD10). The dark periods are represented as gray shaded areas. (D) Plots of locomotor activity onset (black), activity offset (gray), GCaMP onset (green), GCaMP offset (light green), and GCaMP peak (magenta) of mean ± SEM (left column) and individual (right column) in control and *Avp-CK1δ^-/-^* mice. Identical marker shapes indicate data from the same animal. n = 6 for control, n = 4 for *Avp-CK1δ^-/-^*.

**Figure 7.**
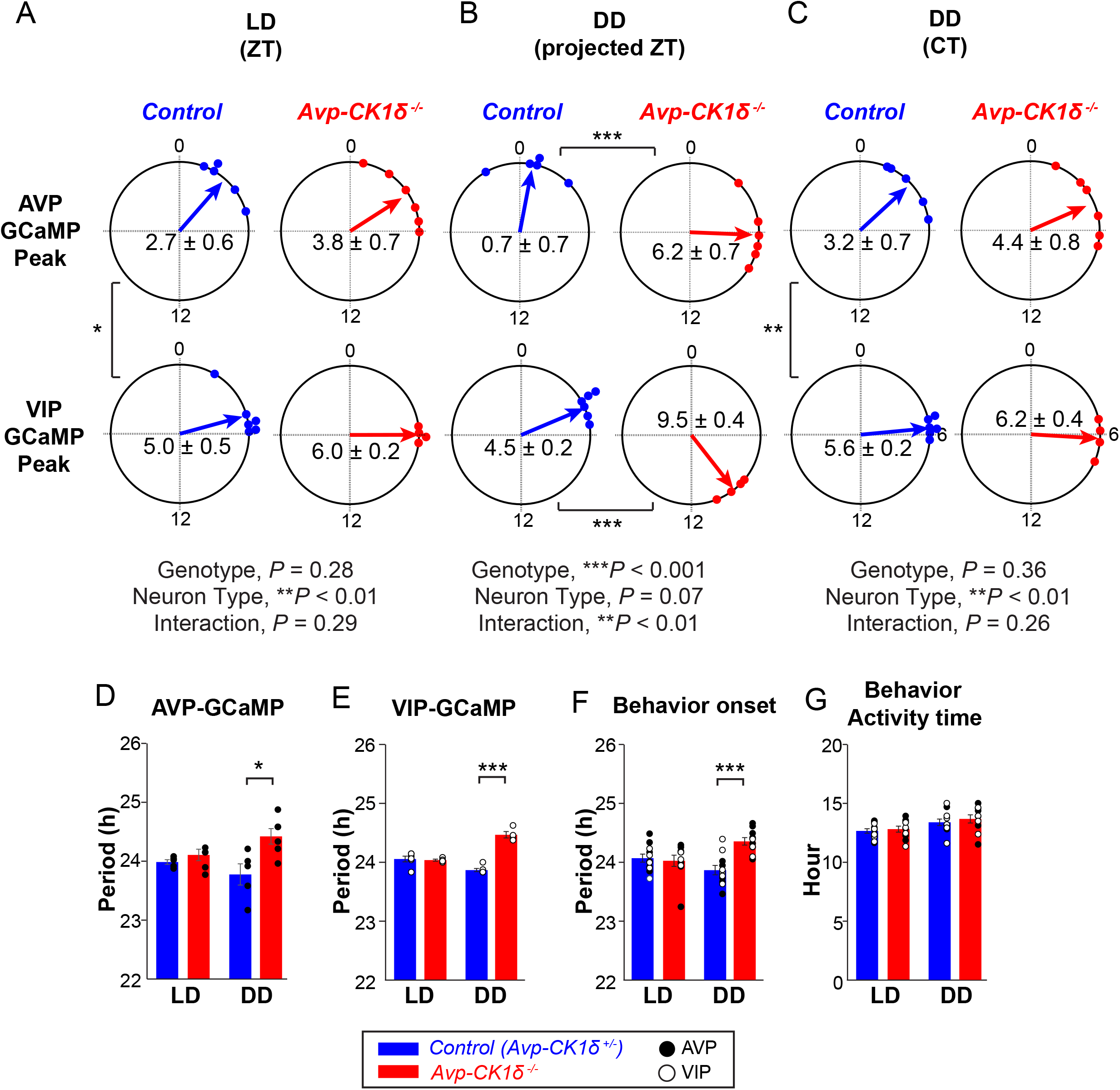
Periods of AVP-neuronal, VIP-neuronal, and behavior rhythms are similarly lengthened in *Avp-CK1δ^-/-^* mice in vivo. (A-C) Peak phases of AVP and VlP GCaMP fluorescence rhythms in LD (LD1-5, A) or DD in projected ZT (DD8-10, B) or CT (C) were shown as Rayleigh plots. Individual dots indicate the peak phases of each mouse. (D) Periods of AVP-neuronal GCaMP fluorescence rhythm in LD (LD1-5, left) or DD (DD8-10, right). (E) Periods of VIP-neuronal GCaMP fluorescence rhythm in LD or DD. (F) Periods of locomotor activity onset in LD or DD. Black circle, data from AVP-GCaMP experiment; white circle, data from VIP-GCaMP experiment. Data from AVP-GCaMP and VIP-GCaMP experiments were combined for statistical analysis. (G) Activity time of locomotor activity rhythm in LD (LD1-5, left) or in DD (DD8-10, right). Blue, Control (*Avp-CK1δ^+/-^* i.e., *Avp-Cre; CK1δ^wt/flox^);* red, *Avp-CK1δ^-/-^*. Values are mean ± SEM. n = 5 for AVP-GCaMP:Control, n = 6 for *AVP-GCaMP: Avp-CK1δ^-/-^* n = 6 for VlP-GCaMP:Control, n = 4 for VIP-GCaMP:*Avp-CK1δ^-/-^*. *\*P* < 0.05; \*\**P* < 0.01; \*\*\**P* < 0.001 by Harrison-Kanji test followed by Watson-Williams test (A to C), or by two-tailed Welch’ s *t*-test (D to G). *P* values of Rayleigh test were <0.01 for all circular data.

Overall, the peak phases of GCaMP signals in LD were similar between control and *Avp-CK1δ^-/-^* mice for both AVP and VIP neurons (AVP: control, ZT 2.74 ± 0.57; *Avp-CK1δ^-/-^* ZT 3.82 ± 0.74, Fig. 7A top; VIP: control, ZT 4.98 ± 0.54; *Avp-CK1δ^-/-^*, ZT 5.99 ± 0.17, Fig. 7A bottom). AVP neurons peaked earlier than VIP neurons (control, *P* < 0.05; *Avp-CK1δ^-/-^*, *P* = 0.07) in LD. These peak phase relationships were essentially maintained even in DD. Due to the lengthening of the free-running period, the peak phase of AVP-neuronal [Ca^2+^]_i_ in projected ZT was significantly later in *Avp-CK1δ^-/-^* during day 8-10 in DD (control, projected ZT 0.72 ± 0.69; *Avp-CK1δ^-/-^*, projected ZT 6.15 ± 0.65, *P* < 0.001, Fig. 7B top), but not different in CT (circadian time) defined by the onset of locomotor activity as CT12 (control, CT 3.16 ± 0.66; *Avp-CK1δ^-/-^*, CT 4.43 ± 0.76, Fig. 7C top). The situation of VIP-neuronal [Ca^2+^]_i_ in DD was similar (control, projected ZT 4.48 ± 0.22; *Avp-CK1δ^-/-^*, projected ZT 9.47 ± 0.36, *P* < 0.001, Fig. 7B bottom; control, CT 5.64 ± 0.17; *Avp-CK1δ^-/-^*, CT 6.23 ± 0.44, Fig. 7C bottom). Thus, the phase relationship between [Ca^2+^]_i_ rhythms of AVP and VIP neurons remained unaltered in *Avp-CK1δ^-/-^* mice. Note that the cellular circadian period was artificially lengthened only in AVP but not VIP neurons. These results suggested that AVP neurons can entrain VIP neurons in vivo.

## Discussion

In the present study, we showed that the period lengthening of the behavior rhythm caused by pan-SCN deletion of *CK1δ* was comparable with that by AVP neuron-specific deletion. In contrast, we did not see a change in the behavioral period in mice with VIP neuron-specific *CK1δ* deficiency. PER2::LUC rhythms in slices did not elucidate the presumed lengthening of the ensemble period of SCN network in *Avp-CK1δ^-/-^* mice. On the other hand, in vivo [Ca^2+^]_i_ rhythms in both SCN AVP and VIP neurons coherently oscillated in *Avp-CK1δ^-/-^* mice with a lengthened period along with the behavior rhythm. Collectively, these data suggest that AVP neurons function as the primary circadian pacesetter in vivo to determine the ensemble period of the SCN network.

### AVP neurons are the principal determinant of the ensemble period of SCN in vivo

Previous studies utilizing neuron type-specific genetic manipulations of cellular periods have suggested the importance of NMS neurons, AVP neurons, Drd1a neurons, and VPAC2 neurons in setting the ensemble period of the SCN (*22, 24, 25, 34, 42*). In contrast, VIP neurons may not contribute significantly to the pace-setting of the SCN network, although VIP is a critical synchronizer for SCN neurons.

Recently, Hamnett et al. demonstrated that lengthening cellular periods in VPAC2 (VIPR2)-expressing neurons also lengthens the free-running period of circadian wheel-running rhythm (*25*). However, PER2::LUC rhythms of SCN slices did not reflect the behavioral period of these mice. Such inconsistency between in vivo and ex vivo periods resembles our current and previous observations for AVP neuron-specific lengthening of cellular periods. Because VPAC2 neurons include ~85% of AVP neurons, their and our findings implicate that the ability of VPAC2/AVP neurons to set the ensemble period of the SCN network is less potent ex vivo than in vivo. Another study by the same group suggested that VPAC2/AVP neurons additionally require the contribution of VIP neurons to dictate the SCN slice period (*28*). This view also explains that the cellular period lengthening in NMS neurons, including both AVP and VIP neurons, successfully lengthened the periods of both SCN slices and behavior (*34*). On the other hand, an essential significance of this study is the demonstration that in vivo AVP neurons alone can entrain other SCN neurons, such as VIP neurons, and determine the SCN ensemble period to control behavior rhythm.

### The stability of the SCN circadian period differs between ex vivo and in vivo

So, what made the difference in periods between in vivo and ex vivo? AVP neurons may require direct or indirect connections with extra-SCN regions, lost in slices, to exert influence on the entire SCN. Another possibility, which seems more appealing to us, is that slicing damaged the usual connections and humoral environment inside the SCN so that AVP neurons are unable to transmit period information to other SCN neurons. The conservation of phase relationship between [Ca^2+^]_i_ rhythms of AVP and VIP neurons in vivo in *Avp-CK1δ^-/-^* mice may prefer the latter possibility. We previously showed that GABA from AVP neurons is unnecessary to transmit their cellular period to the behavioral period (*36*). Therefore, the peptidergic transmission of SCN AVP neurons, such as those mediated by AVP, NMS, cholecystokinin, and Prokineticin 2, may be responsible for the ensemble period setting. In general, peptides are engaged in volume transmission rather than synaptic transmission (*43*). Therefore, the peptidergic transmission by AVP neurons might be more diffusible and attenuated in slice cultures. Also, because AVP neurons are distributed around the core, the ratio of AVP neurons to core cells was smaller in coronal sections containing both shell and core than in vivo. Such a condition may be another reason why the cellular period of AVP neurons was not reflected throughout the SCN in slices.

In this context, it was surprising that PER2::LUC rhythms were dissociated between shell and core in the SCN slices of *CaMKIIα-CK1δ^-/-^* mice. Because, in contrast to *Avp-CK1δ^-/-^* mice, all SCN neurons were supposed to lack *CK1δ*. A simple explanation for this observation is that *CK1δ* contributes to the cellular period setting differently between shell and core cells. Its deficiency might lengthen the cellular period more in the shell than in the core, resulting in chimerism of the cellular periods within the SCN, as in *Avp-CK1δ^-/-^* mice. It is technically challenging to precisely measure the intrinsic cellular period of each type of *CK1δ^-/-^* SCN neurons (e.g., AVP neurons and VIP neurons) under conditions in which extracellular inputs are completely excluded (*10, 11, 44*). The initial periods of PER2::LUC rhythms in the culture might reflect but were not the same as the intrinsic cellular periods. This is because there should be some intercellular communications that could affect the cellular periods. Indeed, the phases of PER2::LUC rhythms in the core cells were relatively advanced in *Avp-CK1δ^-/-^* mice compared to those in control mice (Fig. 4D-F). This observation may suggest repulsive interaction between the shell and the core in chimeric *Avp-CK1δ^-/-^* mice.

The lengthening of the behavioral periods by 0.8~0.9 h in *Avp-CK1δ^-/-^* and *CaMKIIα-CK1δ^-/-^* mice seemed to be less than the 1.5~2 h lengthening observed in *CK1δ^-/-^* peripheral cells, such as primary mouse embryonic fibroblasts (MEFs) and liver explants (*8*). Unfortunately, we have no information concerning the behavioral free-running period of *CK1δ^-/-^* mice, which died perinatally (*8*). Nevertheless, such an apparent reduction of period lengthening in behavior rhythm may be attributed to the nature of in vivo SCN network. Indeed, the circadian-period alteration caused by a series of CRY1 mutations was much smaller in behavior rhythms than in cellular oscillations of MEFs, converging close to 24 h in behavior (*45*).

In any case, this study demonstrated the importance of measuring the dynamics of the SCN network in vivo. Although the slice preparations are beneficial for examining the spatiotemporal organization of cellular clocks in the SCN, there may still be a gap between in vivo and ex vivo SCNs. Indeed, inconsistencies between the behavioral free-running period and the period of *Per1-Luc, Per1-Gfp*, or PER2::LUC oscillations in the SCN slices have also been reported by previous studies (*22, 46–49*). By combining genetically modified mice and AAV vectors utilizing Cre/loxP and Tet systems, we successfully measured the in vivo cellular oscillations of AVP and VIP neurons separately in mice with AVP neuron-specific gene knockout. Such a dual-targeting strategy would be potent for studying the interaction between two different types of cells within the SCN, a small but complicated network consisting of multiple types of neurons and glial cells (*40*).

### Shapes of calcium rhythm differ between AVP and VIP neurons in the SCN

In vivo calcium activity of AVP neurons was close to the sine curve (Fig. 5C). In contrast, the calcium signal of VIP neurons was close to on and off square pulses (Fig. 6C), which nearly delineated the (subjective) day. Such daily patterns of [Ca^2+^]_i_ were consistent with the previous reports (*18, 36, 41*). In single SCN neurons in culture, clock gene expression rhythms are quasi-sinusoidal, whereas the neuronal activity rhythms are often quasi-rectangular in shape. [Ca^2+^]_i_ rhythms may be the intermediate and variable, which seems to be regulated by both the intercellular neuronal network and the intracellular modulators linking the TTFL to [Ca^2+^]_i_ (*11, 12, 44, 50–52*).

By analogy, AVP-neuronal [Ca^2+^]_i_ rhythm may be regulated primarily by the intracellular TTFL mechanism, while neuronal firing may contribute more significantly to VIP-neuronal [Ca^2+^]_i_. This difference may be related to the seemingly contradictory results that optogenetically increasing the firing rate of VIP neurons in SCN slices caused a phase shift in the PER2::LUC rhythm, but the similar stimulation of recipient VPAC2 neurons did not (*28*). As discussed, some mechanisms in addition to the neuronal activity and Ca^2+^ influx, such as cAMP responsive element (CRE)-dependent transcription, may be essential for modulating TTFL in VPAC2/AVP neurons. The more rectangular Ca^2+^ rhythm in VIP neurons is like an on-and-off switch. This shape may be suitable for setting time frames according to the LD cycle for locomotor activity, sleep-wakefulness, and other bodily functions. Also, it may be beneficial to mediate the light entrainment (*18, 53–55*).

### AVP neurons as the primary oscillating part of the SCN network

As discussed above, AVP neurons are likely the principal pacesetter of the SCN network clock. We previously demonstrated that disrupting cellular clocks by deleting *Bmal1* specifically in AVP neurons drastically attenuated the coherency of circadian behavior rhythm (*Avp-Bmal1^-/-^* mice). It has also been reported that *Bmal1* function in AVP neurons, not VIP neurons, is essential for autonomous network synchrony of the SCN network in slices (*56*). Only a tiny number of *Avp-Bmall^-/-^* mice showed arrhythmicity when home-cage activity was recorded (*22*). However, the same mice exhibit more severe disturbance of wheel-running behavior rhythm resembling those described as “arrhythmic” or “split” in VPAC2 neuron-specific *Bmal1* deletion (*25*). In addition, impaired circadian rhythms of home-cage activity and body temperature in *Avp-Bmall^-/-^* mice are strikingly similar to those of *Nms-Bmal1^-/-^* mice (*35*). The latter has been reported to be arrhythmic in wheel-running rhythm (*34*).

We propose the following model to integrate findings in the previous reports and this study. AVP neurons function as the primary oscillatory part of the SCN network, determining the period and taking charge of the oscillation itself. The cellular clocks of AVP neurons may be sloppy and not necessarily self-sustaining. The sustained oscillation of the cellular clocks of AVP neurons may need to be driven by other SCN neurons. VIP neurons likely play this role, synchronize many AVP neurons to a certain range, and transmit information about external light to regulate AVP neurons’ phases. On the other hand, the cellular clocks of VIP neurons are entrained by AVP neurons and likely be dispensable for circadian pacemaking. Thus, the functional differentiation and reciprocal interaction between cell types in the SCN network exerts the central clock function.

## Materials and Methods

### Animals

*Avp-Cre* and *Vip-tTA* mice were reported previously (*22, 40*). This *Avp-Cre* line is a transgenic mouse harboring a modified BAC transgene, which has an insertion of codon-improved Cre recombinase gene immediately 5’ to the translation initiation codon of exogenous *Avp* gene in the BAC, but without manipulation of the endogenous *Avp* loci in the mouse. Thus, the strain is different from the widely-used *Avp-ires2-Cre* (JAX #023530) that shows hypomorphic expression of AVP (*57*), as well as from another *Avp-ires-Cre* line (*58*). *CK1δ^flox^* (JAX #010487) (*8*), *Bmal1^flox^* mice (JAX #007668) (*59*), and *Vip-ires-Cre* (*Vip^tm1(cre)Zjh^*/J, JAX #010908) were obtained from Jackson Laboratory. *CaMKIIα-Cre* (*Camk2a::iCreBAC*) (*30*) was obtained from the European Mouse Mutant Archive (EMMA #01153). The *Per2::Luc* reporter mice were provided by Dr. Joseph Takahashi (*60*). All lines were congenic on C57BL/6J. *Avp-Cre, CaMKIIα-Cre, Vip-ires-Cre*, and *Vip-tTA* mice were used in hemizygous or heterozygous condition. Mice were maintained under a strict 12 h light/12 h dark cycle in a temperature- and humidity-controlled room and fed *ad libitum*. All experimental procedures involving animals were approved by the appropriate institutional animal care and use committees of Kanazawa University and Meiji University.

### Behavioral analyses

Male and female *CaMKIIα-CK1δ^-/-^* (*CaMKIIα-Cre; CK1δ^flox/flox^* and *Vip-CK1δ*^-/-^ (*Vip-ires-Cre; CK1δ^flox/flox^*) mice, aged 8 to 20 weeks, were housed individually in a cage placed in a light-tight chamber (light intensity was approximately 100 lux). Spontaneous locomotor activity was monitored by infrared motion sensors (O’Hara) in 1-min bins as described previously (*22*). *CK1δ^flox/flox^* and *CaMKIIa-Cre; CK1δ^wt/flox^* littermates were used together as the control for *CaMKIIα-CK1δ^-/-^* mice. *CK1δ^flox/flox^* and *Vip-ires-Cre; CK1δ^wt/flox^* littermates were used as the control for *Vip-CK1δ*^-/-^ mice. Because two control lines (*CK1δ^flox/flox^* and *Cre; CK1δ^wt/flox^*) for each conditional knockout behaved similarly, we regarded them together as control. Actogram, activity profile, and χ^2^ periodogram analyses were performed via ClockLab (Actimetrics). The free-running period was measured by periodogram for days 10-24 in DD. The activity time was calculated from the daily activity profile (average pattern of activity) of the same 15 days using the mean activity level as a threshold for detecting the onset and the offset of activity time (*22*). Wheel running activity was measured for *Avp-CK1δ^-/-^* (*Avp-Cre; CK1A^flox/flox^*), *Avp-Bmal1^-/-^ (Avp-Cre; Bmal1^flox/flox^*), and control mice as described previously (*61*). Briefly, each mouse was housed in a separate cage equipped with a running wheel (12-cm diameter; SANKO, Osaka, Japan). The cages were placed in ventilated boxes; the light intensity at the bottom of the cage was 200–300 lux. The number of wheel revolutions was counted in 1-min bins. A chronobiology kit (Stanford Software Systems) and ClockLab software (Actimetrics) were used for data collection and analyses.

### Bioluminescence Imaging

The *Avp-CK1δ^-/-^. CaMKIIa-CK1δ^-/-^*, and *Vip-CK1δ^-/-^* mice were further mated with *Per2::Luc* reporter mice (*60*) (*Avp-Cre; CK1δ^flox/flox^*; *Per2::Luc, CaMKIIα-Cre; CK1δ^flox/flox^; Per2::Luc, Vip-Cre; CK1δ^flox/flox^; Per2::Luc*) and compared to control mice (*CK1δ^flox/flox^; Per2::Luc*). Male and female mice aged 8 to 17 weeks were housed in LD before sampling. Coronal SCN slices of 150 μm were made at ZT8– with a linear-slicer (NLS-MT; Dosaka-EM). The SCN tissue at the mid-rostrocaudal region was cultured and its PER2::LUC bioluminescence was imaged every 30 min with an exposure time of 25 min with an EMCCD camera (Andor, iXon3), as described previously (*23*).

Images were analyzed using ImageJ and MATLAB (MathWorks). The resolution of images was adjusted to 4.6 μm/pixel for further analyses. Bioluminescence values in individual pixels were detrended by subtracting 24-h moving average values and then were smoothened with a fifteen-point moving average method. The middle of the time points crossing the value 0 upward and downward was defined as peak phases, whereas the maximum value between these two time points was defined as amplitude. The intervals between two adjacent peak phases were calculated as the periods (*22*). The values of the regions lateral to the SCN were regarded as background. Therefore, pixels with values lower than the background were eliminated for further analyses. Periods of individual pixels’ oscillation were calculated for every cycle. Then, square regions of interest (ROI, 15 x 15 pixels) were defined in the shell and core of the left and right SCN. Pixels with a period shorter than 4 h or longer than 40 h were eliminated to calculate the mean period, amplitude, and peak phase. Pixels in the left and right ROIs were first processed separately. Then the averaged values of these two were considered representatives of individual mice.

### Viral vector and Surgery

The AAV-2 ITR containing plasmids *pGP-AAV-CAG-FLEX-jGCaMP7s-WPRE* (Addgene plasmid #104495, a gift from Dr. Douglas Kim & GENIE Project) (*37*) and *pAAV-TRE-EGFP* (Addgene plasmid #89875, a gift from Dr. Hyungbae Kwon) (*62*) were obtained from Addgene. *pAAV-TRE-jGCaMP7s* was described previously (*40*). Recombinant AAV vectors (AAV2-rh10) expressing *jGCaMP7s (AAV-CAG-DIO-jGCaMP7s, AAV-TRE-jGCaMP7s*)in a Cre-dependent manner or tTA-dependent manner were produced using a triple-transfection, helper-free method and purified as described previously (*22*). The titers of recombinant AAV vectors were determined by quantitative PCR: *AAV-CAG-DIO-jGCaMP7s*, 3.4 x 10^13^; AAV-*TRE-jGCaMP7s*, 6.3 x 10^11^; and *AAV-TRE-EGFP*, 5.7 x 10^11^ genome copies/ml. Stereotaxic injection of AAV vectors was performed as described previously (*22*). Two weeks after surgery, we began monitoring the mice for their locomotor activity.

### In vivo fiber photometry

We used six *Avp-CK1δ^-/-^* x *Vip-tTA (Avp-Cre; CK1δ^flox/flox^; Vip-tTA*) mice, four *Avp-CK1δ^-/-^ (Avp-Cre; CK1δ^flox/flox^*) mice, and 11 control mice (eight *Avp-Cre; CK1δ^wt/flox^; Vip-tTA* and three *Avp-Cre; CK1δ^wt/flox^*). We combined the data of *Avp-Cre; CK1δ^flox/flox^; Vip-tTA* and *Avp-Cre; CK1δ^flox/flox^* for AVP neurons recording considering the lack of inter-group differences. Additional four mice (*Vip-tTA*) were used for control experiments to measure EGFP signal in VIP neurons. The mice were anesthetized by administering a cocktail of medetomidine (0.3 mg/kg), midazolam (4 mg/kg), and butorphanol (5 mg/kg) and were secured at the stereotaxic apparatus (Muromachi Kikai). Lidocaine (8 %) was applied for local anesthesia before making the surgical incision. We drilled small hole in the exposed region of the skull using a dental drill. We injected 0.5-1.0 μL of the virus (*AAV-CAG-DIO-jGCaMP7s* or *AAV-TRE-jGCaMP7s*) (flow rate= 0.1 μL/min) at the right suprachiasmatic nucleus (SCN, posterior: 0.5 mm, lateral: 0.25 mm, depth: 5.7 mm from the bregma) with a 33 G Hamilton Syringe (1701RN Neuros Syringe, Hamilton) to label AVP neurons. Subsequently, we placed an implantable optical fiber (400 μm core, N.A. 0.39, 6 mm, ferrule 2.5 mm, FT400EMT-CANNULA, Thorlabs) above the SCN (posterior: 0.2 mm, lateral: 0.2 mm, depth: 5.2 - 5.4 mm from the bregma) with dental cement (Super-bond C&B, Sun Medical). The dental cement was painted black. Atipamezole (0.3 mg/kg) was administered postoperatively to reduce the anesthetized period. The mice were used for experiments 2–7 weeks after the virus injection and optical fiber implantation. Their ages were 3–10 months old including both male and female.

A fiber photometry system (COME2-FTR, Lucir) was used to record the calcium signal of AVP neurons in freely moving mice (*36, 63*). Fiber-Coupled LED (M470F3, Thorlabs) with LED Driver (LEDD1B, Thorlabs) was used as an excitation blue light source. The light was reflected by a dichroic mirror (495 nm), went through an excitation bandpass filter (472/30 nm), then to the animal via a custom-made patch cord (400 um core, N.A. 0.39, ferrule 2.5 mm, length 50 cm, COME2-FTR/MF-F400, Lucir) and the implanted optical fiber. We detected the jGCaMP7s fluorescence signal by a photomultiplier through the same optical fibers and an emission bandpass filter (520/36 nm); furthermore, we recorded the signal using Power Lab (AD Instruments) with Lab Chart 8 software (AD Instruments). The excitation blue light intensity was 2.5-100 μW at the tip of the patch cord of the animal side. We recorded the same for 30 s every 10 min in two weeks to reduce photobleaching. During the recording, the mouse was housed in a 12-h light–dark cycle for more than 5 days (LD condition), then moved to continuous darkness for approximately 10 days (DD condition) in a custom-made acrylic cage surrounded by a sound-attenuating chamber. A rotary joint for the patch cord was stopped during the recording to prevent artificial baseline fluctuation. The animal’s locomotor activity was monitored using an infrared sensor (Supermex PAT.P and CompACT AMS Ver. 3, Muromachi Kikai).

The detected jGCaMP7s signal was averaged within a 30 s session (*36*). To detrend the gradual decrease of the signal during recording days, ±12 h average from the time (145 points) was calculated as baseline (F). The data was subsequently detrended by the subtraction of F (ΔF). Then, the ΔF/F value was calculated. To determine the peak phase of jGCaMP7s calcium signal, ΔF/F were smoothened with a 21-points moving average, then the middle of the time points crossing value 0 upward and downward were defined as peak phases. Additionally, the intervals between peak phases were defined as the periods (*22*). A double-plotted actogram of jGCaMP7s or EGFP signal was designed by converting all ΔF to positive values by subtracting the minimum value of ΔF. Subsequently, these values were multiplied with 100 or 1000 and rounded-off. The plots were made via ClockLab (Actimetrics) with normalization in each row. A double-plotted actogram of locomotor activity was also prepared and overlaid on that of jGCaMP7s signal.

The onset and offset of locomotor activity were determined using the actogram of locomotor activity. Initially, we attempted to automatically detect the onset and offset; however, it was followed by a manual visual inspection, along with modifications by the experimenter. To calculate circadian time (CT) of the peak phases of GCaMP signal, we have defined the regression line of locomotor activity onsets as CT12.

We confirmed the jGCaMP7s expression and the position of the optical fiber by slicing the brains in 100 μm coronal sections using a cryostat (Leica). The sections were mounted on glass slides with a mounting medium (VECTASHIELD HardSet with DAPI, H-1500, Vector Laboratories) and observed via epifluorescence microscope (KEYENCE, BZ-9000E).

### Statistical analysis

All results are expressed as mean ± SEM. For comparisons of two groups, two-tailed Student’s or Welch’s t tests were performed. For comparisons of multiple groups with no difference of variance by Bartlett test, three-way repeated measures ANOVA followed by post-hoc Ryan’s test were performed. For comparisons of multiple groups with difference of variance by Bartlett test, nonparametric tests, Kruskal-Wallis test with post-hoc Mann-Whitney U test with Bonferroni correction, Friedman rank sum test followed by Wilcoxon signed rank test with Bonferroni correction, and Wilcoxon signed rank test were performed with EZR, a graphical user interface for R (*64*). For circular data, Rayleigh test, Watson-Williams test, and Harrison-Kanji test were performed with Circstat MATLAB Toolbox for Circular Statistics (*65*). All *P* values less than 0.05 were considered as statistically significant. Only relevant information from the statistical analysis was indicated in the text and figures.

## Acknowledgements

We thank D.R. Weaver for the *CK1δ^flox^* mouse; G. Schütz for the *CaMKIIα-Cre (Camk2a::iCreBAC*) mouse; Z. J. Huang for the *Vip-ires-Cre* mouse; J. Takahashi for the *Per2::Luc* mouse; C.J. Weitz for the *Bmal1δ_flox_* mouse; H. Okamoto for technical support to generate *Avp-Cre* mouse; Penn Vector Core for *pAAV2-rh10;* D. Kim & GENIE Project for *pGP-AAV-CAG-FLEX-jGCaMP7s-WPRE*; and H. Kwon for *pAAV-TRE-EGFP*. We thank all lab members, including A. Matsui, M. Fukushi, M. Kawabata, M.T. Islam, and Y. Nishiwaki.

## Author contributions

Conceptualization: YT, MM

Methodology: YT, YP, TD, SH, KY, TM, MM

Investigation: YT, MM Visualization: YT, MM Supervision: YT, MM

Writing—original draft: YT, MM

Writing—review & editing: YT, YP, TD, SH, KY, TM, MM

## Data and materials availability

All data are available in the main text or the supplementary materials.

