## Supplemental Figures for "AVP neurons act as the primary circadian pacesetter cells in vivo"

Sup. Figure S1

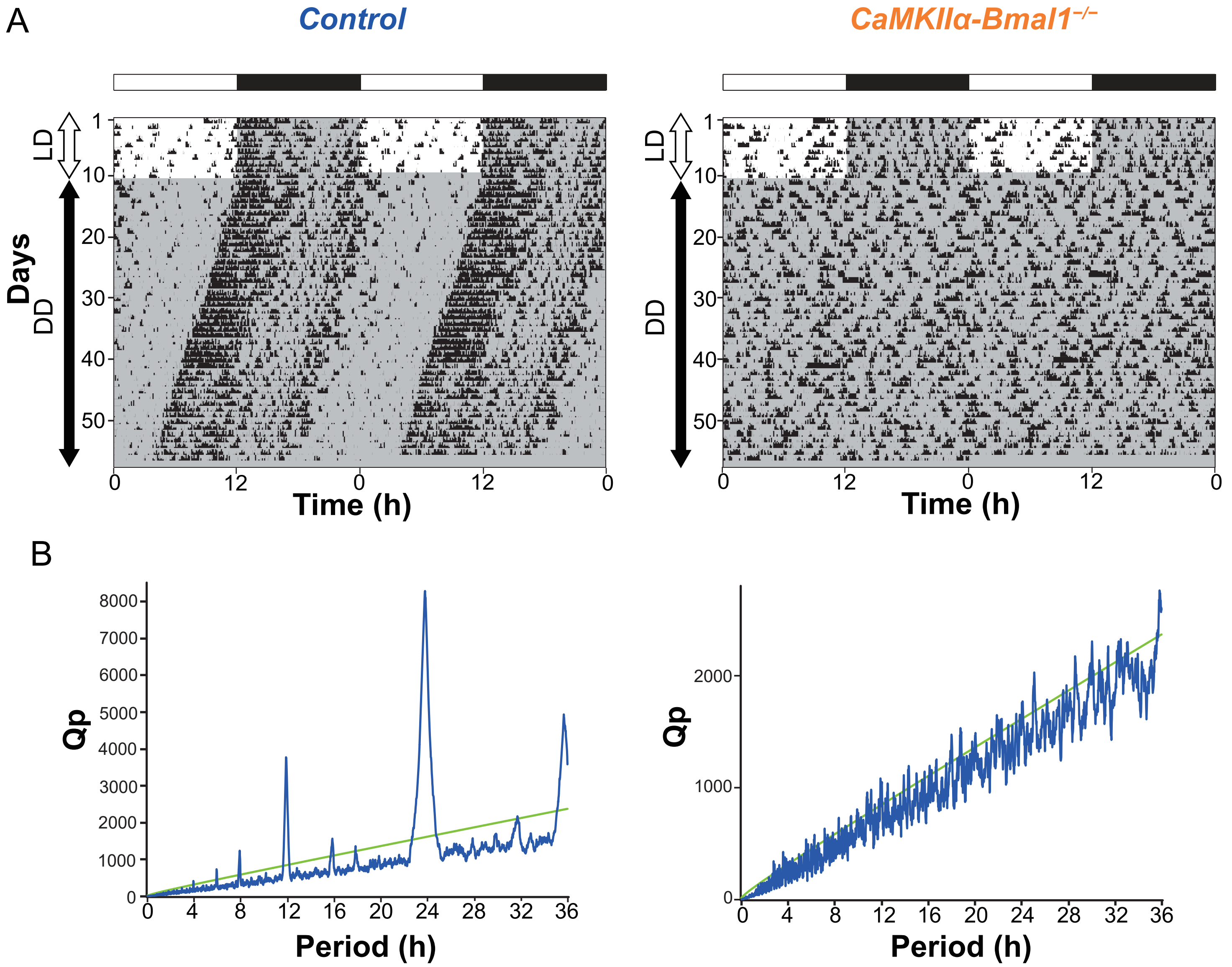


Sup. Figure S1. *CaMKIIα-Bmal1*$^{-/-}$ mice are arrhythmic in DD.
A. Representative locomotor activity of control and *CaMKIIα-Bmal1*$^{-/-}$ mice. Gray shading indicates the time when lights were off.
B. Representative periodograms of the locomotor activity rhtyhms of control (left) and *CaMKIIα-Bmal1*$^{-/-}$ mice (right) in DD.

Sup. Figure S2

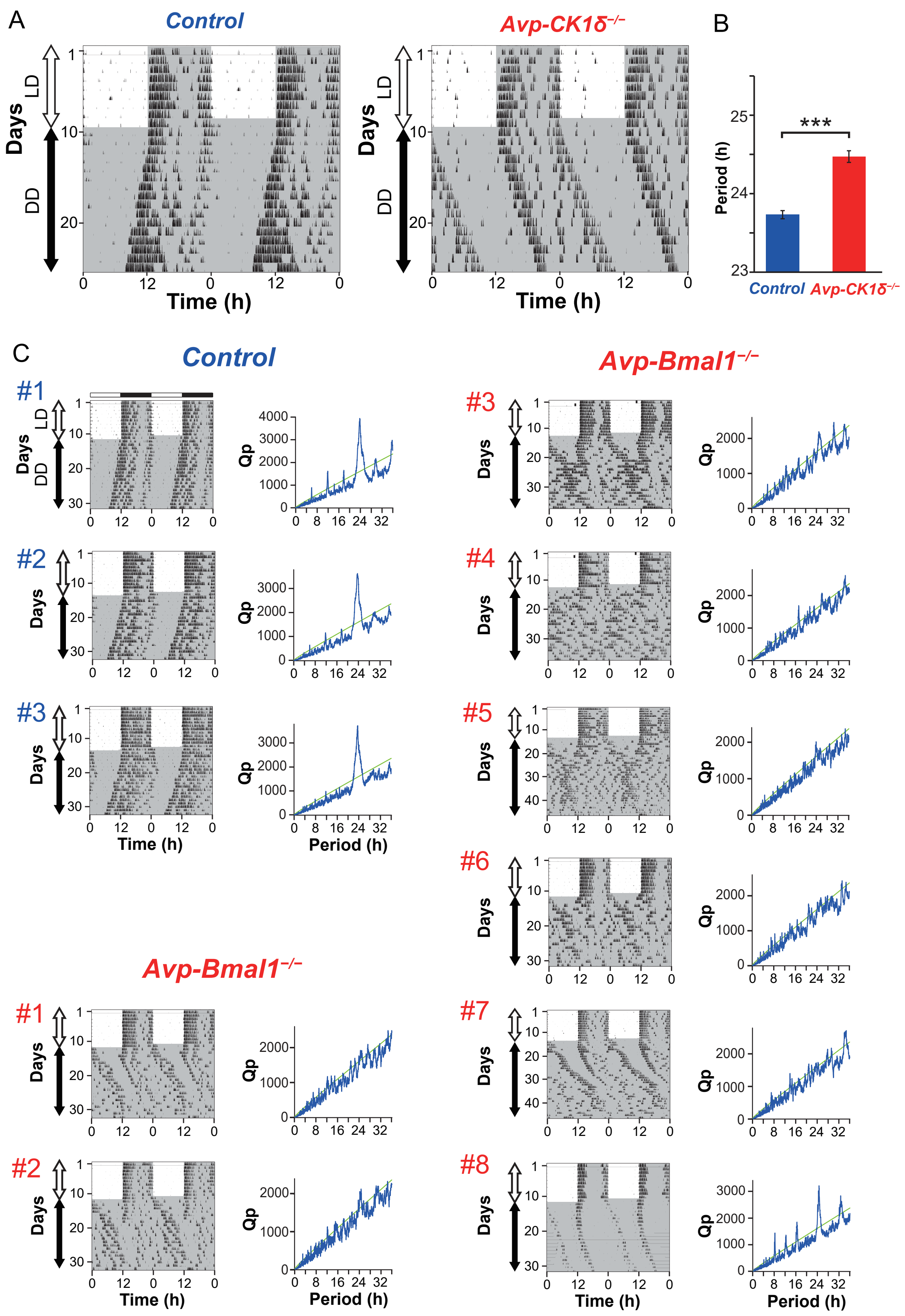


Sup. Figure S2. Re-evaluation of circadian behavior rhythms in *Avp-CK1δ*$^{-/-}$ and *Avp-Bmal1*$^{-/-}$ mice by wheel-running activity.
(A) Representative wheel-running activity of control and *Avp-CK1δ*$^{-/-}$ mice. Gray shading indicates the time when lights were off.
(B) The free-running period of wheel-running activity in DD.
Values are mean ± SEM; n = 4 for control, n = 5 for *Avp-CK1δ*$^{-/-}$ mice. ***$P < 0.001$ by two-tailed Student' s t test.
(C) Left: Actograms of the wheel-running activity of three control and eight *Avp-Bmal1*$^{-/-}$ mice. Gray shading indicates the time when lights were off. Right: Periodograms of the individual wheel-running activity rhythms in the last seven days in DD. Most *Avp-Bmal1*$^{-/-}$ mice were arrhythmic.

### Sup. Figure S3

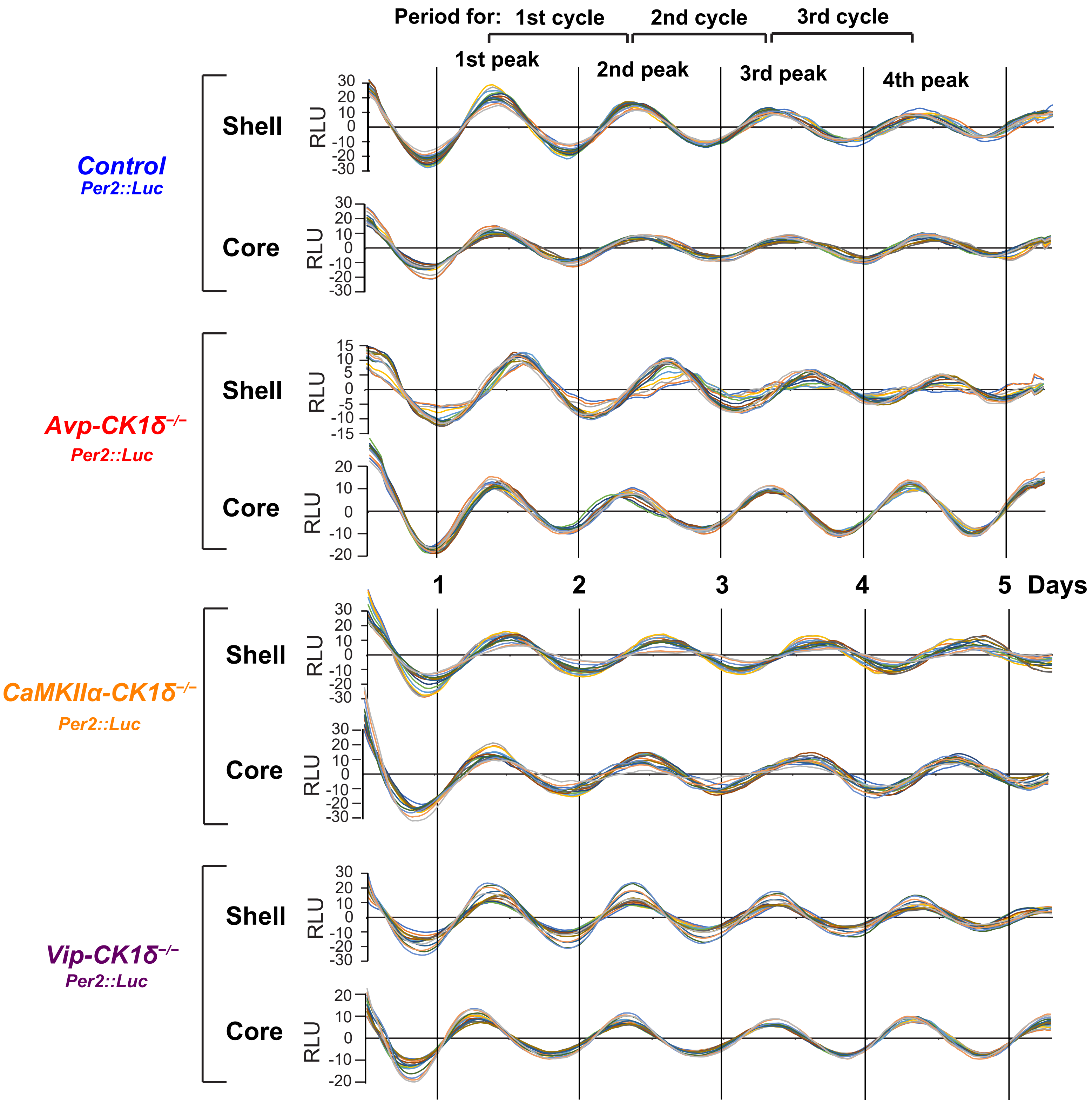


Sup. Figure S3. Single-pixel trajectories of Per2::LUC oscillation in the SCN slices of *CK1δ* conditional knock-out mice. Representative bioluminescence data of 15 pixels in a row along the mediolateral axis within ROIs (15 x 15 pixels) are shown for each region and mouse line. All data were detrended, smoothed, and aligned at ZT12 on the day of slicing as starting points. Black vertical lines labeled day 1 indicate projected ZT0. These data were used for statistical analysis in Figure 4.

### Sup. Figure S4

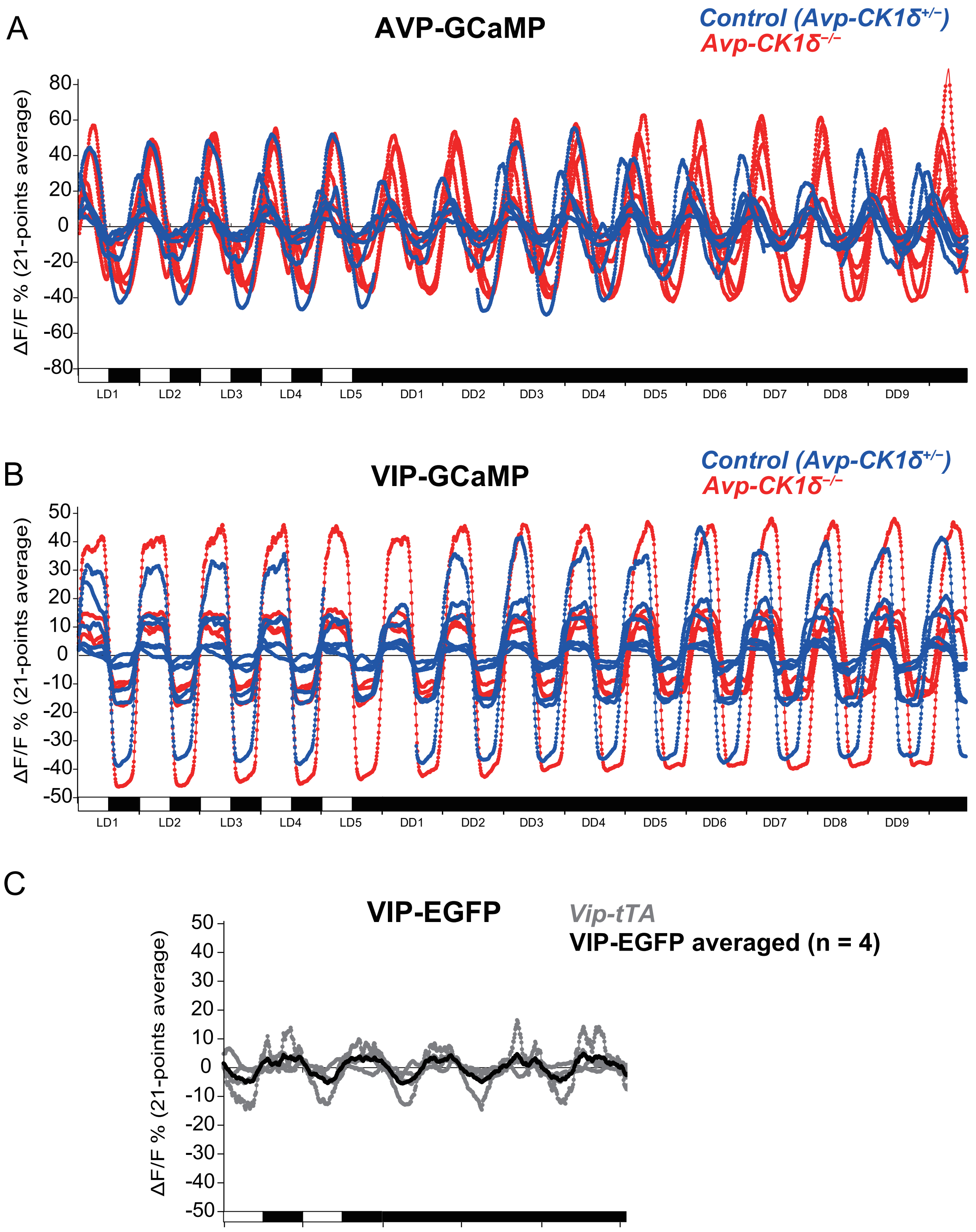


Sup. Figure S4. Individual trajectories of GCaMP and EGFP fluorescence in SCN neurons in vivo.
(A, B) Continuous recordings of GCaMP fluorescence from SCN AVP neurons (A) or VIP neurons (B) for 15 days (5 days in LD, 10 days in DD). Red, *Avp-CK1δ*$^{-/-}$ (n = 6, 4); blue, control (*Avp-CK1δ*$^{+/-}$, i.e., *Avp-Cre; CK1δ*$^{wt/flox}$, n = 5, 6).
(C) Continuous recordings of EGFP fluorescence from SCN VIP neurons for 5 days (2 days in LD, 3 days in DD). Gray, individual trace; black, average.
